# A novel post-developmental role of the Hox genes underlies normal adult behaviour

**DOI:** 10.1101/2021.08.29.458086

**Authors:** A. Raouf Issa, Claudio R. Alonso

## Abstract

The molecular mechanisms underlying the stability of mature neurons and neural circuits are poorly understood. Here we explore this problem and discover that the *Hox* genes are a component of the genetic programme that maintains normal neural function in adult *Drosophila*. We show that post-developmental downregulation of the *Hox* gene *Ultrabithorax* (*Ubx*) in adult neurons leads to substantial anomalies in flight. Mapping the cellular basis of these effects reveals that *Ubx* is required within a subset of dopaminergic neurons, and cell circuitry analyses and optogenetics allow us to link these dopaminergic neurons to flight control. Functional imaging experiments show that *Ubx* is necessary for normal dopaminergic activity, and neuron-specific RNA-sequencing defines two previously uncharacterised ion channel-encoding genes as potential mediators of *Ubx* behavioural roles. Our study thus reveals a novel role of the *Hox* system in controlling adult behaviour and neural function.

## MAIN TEXT

At the end of the developmental process, individual neurons must deactivate the genetic programmes that guided their patterning and differentiation, and switch on molecular programmes that ensure neural maintenance and physiology. While considerable efforts have been invested into the decoding of neural developmental programmes, less is known about the identity and influence of genetic systems in the fully mature, post-mitotic neuron of the adult organism. Here we investigate this problem in the fruit fly, *Drosophila melanogaster*, taking advantage of the genetic accessibility and detailed understanding of neural processes in this organism.

Recent observations in our laboratory suggested that over-expression of particular transcription factors in post-mitotic neurons had impact on neural physiology (*1–4*). Amongst these factors, are the *Hox* genes, which encode a family of developmental factors whose activities control cell differentiation programs in animals as diverse as insects and mammals (*5–7*). Hox gene expression patterns have been studied in great detail during embryogenesis (*8, 9*) but recent data from our laboratory and elsewhere (*3, 10*) show that *Hox* genes are also expressed in adult forms (Figure 1A), yet, the biological roles of this late phase of expression are not known. We hypothesised that the *Hox* genes may contribute to the genetic programme underlying brain stability and used the *Drosophila* adult as a system to explore this notion, seeking to establish the biological roles of adult post-mitotic neural *Hox* gene expression.

**Fig. 1.**
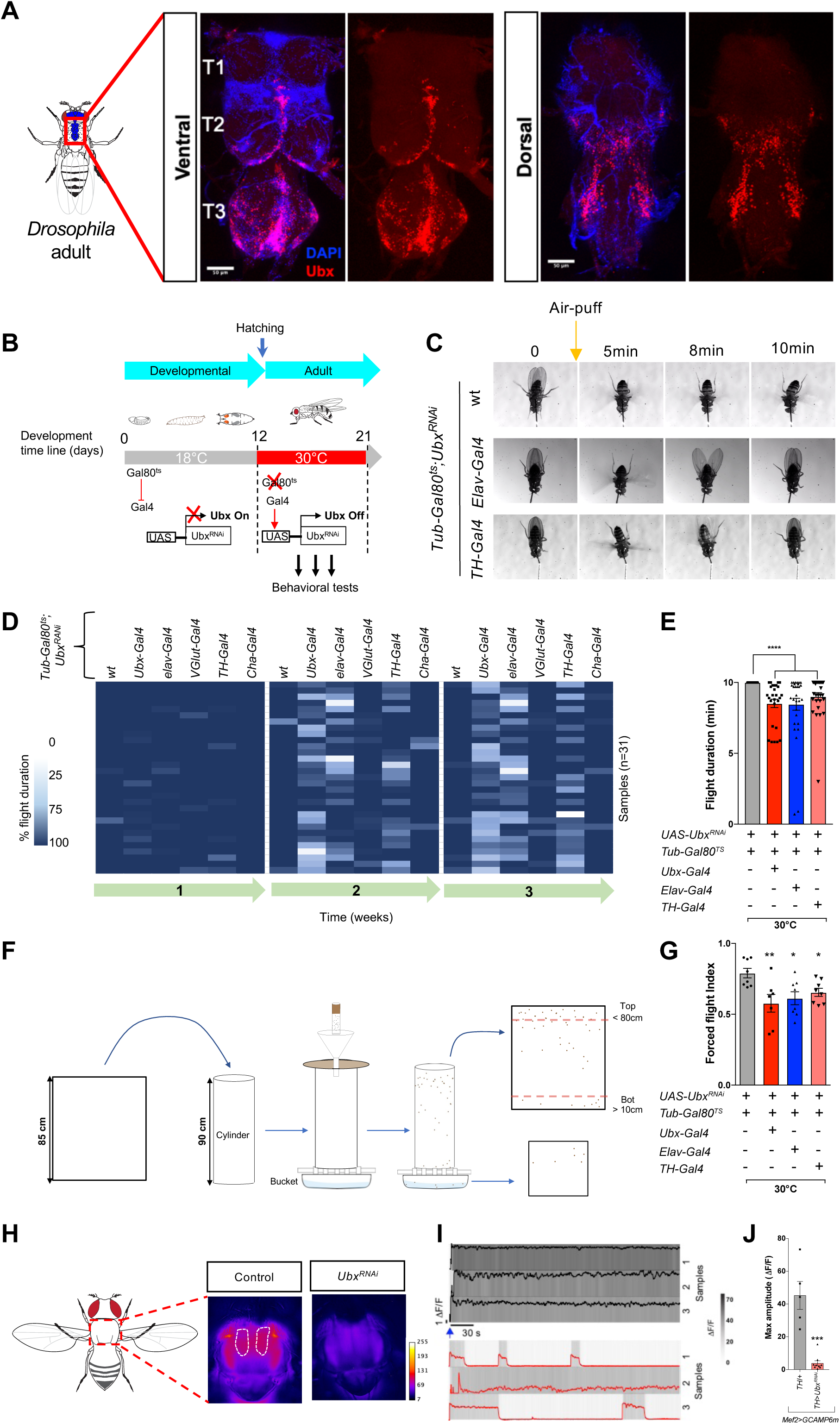
*Ubx* is necessary for normal flight maintenance in adult *Drosophila*. (**A**) Confocal images of the whole ventral nerve cord (VNC) of adult *Drosophila* showing expression of Ubx protein in specific regions of the adult VNC (T1-T3, indicate the respective thoracic segments). (**B**) Strategy for conditional neuronal downregulation of *Ubx* in the adult. Flies expressing *Ubx^RNAi^* under temperature-sensitive Gal80ts repression, were exposed to elevated temperatures (30°C) upon hatching to allow active downregulation of *Ubx*. At permissive temperature (18°C) Gal80ts protein is functional and represses expression of *Ubx^RNAi^* (‘Ubx ON’); conversely, expression of *Ubx^RNAi^* is activated in cells at temperatures above 30°C (‘Ubx OFF’). Behavioural experiments were performed on adult males and females in Ubx ON/OFF modes. (**C**-**E**) Evaluation of flight performance on tethered *Drosophila*. (**C**) Representative flight duration is indicated by video snapshots of tethered wild-type (wt), pan-neuronal (*Tub-Gal80ts*;*Elav-Gal4*, *Ubx^RNAi^*) and dopaminergic (*Tub-Gal80ts*;*TH-Gal4*, *Ubx^RNAi^*) *Ubx^RNAi^* expressing flies, at 30°C (C). The orange arrow indicates the induction stimulus (air-puff). (**D** and **E**) Flight duration represented as a heat map, and as averages of flight duration at week 2, for flies expressing *Ubx^RNAi^* in different nerve cells (*Ubx/elav/Vglut/TH/Cha>Ubx^RNAi^*) compared to controls (*UAS-Ubx^RNA^*^i^) under Gal80ts repression (week 1= 6-7 days; week 2= 8-10 days; week 3= 14-15 days; n >20) (Supportive movie: Movies 1). (**F**). Cartoon illustrating the forced flight test procedure (for details, see materials and methods). (**G**) Average forced flight index at week 2 of *Ubx>Ubx^RNAi^*, *elav>Ubx^RNAi^*, and *TH>Ubx^RNAi^* flies reared at 30°C compared to controls (*UAS-Ubx^RNA^*^i^) (Each data point represents a group of n>12 flies). (**H**) Representative raw images of GCAMP fluorescence (warmer color) in flight muscles. (**I**) representative calcium signal traces in dorsal longitudinal flight muscle (Dlm) from control (*Mef2^LexA^,GCAMP6m^LexAOP^/Tub-Gal80^ts^;* TH-Gal4/FLP) and experimental (*Mef2^LexA^,GCAMP6m^LexAOP^/Tub-Gal80^ts^;* TH-Gal4/Ubx^RNAi^) flies. (**J**) Normalised mean of GCaMP signals maximum during sustained flight (n = 5-10) )(Supportive movie: Movies 2). All error bars represent SEM. Significant values in all figures based on Mann-Whitney U test or one-way ANOVA with the post hoc Tukey-Kramer test: ^∗^p < 0.05, ^∗∗^p < 0.01, and ^∗∗∗^p < 0.001.

To determine the role of the *Hox* genes in the adult, without affecting their previous developmental functions, we applied a conditional expression strategy that maintains normal *Hox* expression during development and reduces expression at eclosion time (Fig. 1A). For this we used the Gal80/Gal4 system (*11, 12*) to drive tissue-specific RNA interference constructs to reduce the expression of one of the posterior *Drosophila Hox* genes, the gene *Ultrabithorax* (*Ubx)* (*13*) (Fig. S1) exclusively in the adult. Analysis of the behavioural impact of these perturbations during the first three weeks of adult life, when targeting interference to all neurons (*elav-Gal4*) or all cells expressing *Ubx* (*Ubx-Gal4*) results in no detectable changes in general fly locomotion, as assessed through standard climbing assays (Fig. S2A-C); this first result suggested that reduction of *Ubx* function in the adult may not impact the normal physiology of motor circuits underlying adult movement on substrate. In contrast, tethered flight experiments(*14, 15*) reveal that RNAi-mediated knockdown of *Ubx* expression does impair flight maintenance (Fig. 1C-E, Fig. S2D-F, and Movie 1) at two weeks of age: expression of *elav*-driven *UAS-Ubx^RNAi^* leads to a marked deficit in the ability of flies to maintain flight following a single air-puff (NB: similar effects on flight were obtained when using a different RNAi line targeting *Ubx* (Figure S1)). Two other independent methods to assess flight performance, i.e. forced flight(*15*) (Fig. 1F-G, S2G), and take-off (*15, 16*) (Fig. S2H-I and Movies S1 and S2) tests confirm that normal expression of *Ubx* is required for normal behaviour in adult flies, revealing a novel role for the Hox system in the control of behaviour of the fully-formed organism.

Seeking to identify the main neuronal types where *Ubx* is required for normal flight, we tested the impact of *Ubx* reduction within the cholinergic (*Cha-Gal4*), glutamatergic (*VGlut-Gal4*) and dopaminergic (*TH-Gal4*) systems (Fig. 1 and Fig. S2). The results of these perturbations show that reduction of *Ubx* within the dopamine (TH) domain is sufficient to phenocopy the effects observed within the pan-neural domain (*elav-Gal4*) (Fig. 1C-G and Fig. S2D-I); in contrast, no significant effects are detected after *Ubx* reduction in glutamatergic or cholinergic neurons (Fig. 1D, Fig. S2D-K). These observations imply that dopaminergic neurons (here on termed TH neurons) might be particularly sensitive to reductions in *Ubx* input, arguing that normal *Ubx* expression is required for normal function of TH neurons. Furthermore, fluorescence imaging of flight muscle (*15, 17*) expressing a calcium reporter (GCaMP6m) (Fig. 1H-J; and Movies 2 and 3) shows that *TH>Ubx^RNAi^* leads to a significant reduction of flight muscle activity. Altogether, these data indicate that normal expression of *Ubx* in dopaminergic neurons is required for normal flight maintenance in adult flies.

The functional experiments described above implied that *Ubx* might be expressed in the adult dopaminergic system. *Ubx* is expressed in the adult ventral nerve cord (VNC) – the neural equivalent of the mammalian spinal cord (*18*) – across extensive domains on its dorsal and ventral aspects (Fig. 1A, and previous observations (*3, 10*)). To probe specific expression within the dopaminergic system, we labelled the TH domain using nuclear GFP (NLS::GFP) under a dopaminergic driver (*TH-Gal4*) (Fig. 2A), and mapped *Ubx* expression over this territory. The results of this experiment show that Ubx protein was indeed expressed in approximately ∼ 30% (5±0.9) and 60% (17±0.5) of all adult dopaminergic neurons within the dorsal and ventral VNC regions, respectively (Fig. 2B). Furthermore, dopaminergic-specific conditional reduction of *Ubx* expression triggered at adult eclosion time (*TH-Gal4; tub-Gal80^ts^; UAS-Ubx^RNAi^*) resulted in ∼50% downregulation of *Ubx* expression within the TH domain after two weeks of treatment (Fig. 2C-E). These expression analyses confirm that Ubx is normally expressed in adult TH neurons, and that our post-developmental manipulations of *Ubx* expression lead to a substantial reduction in protein formed within the dopaminergic system.

**Fig. 2.**
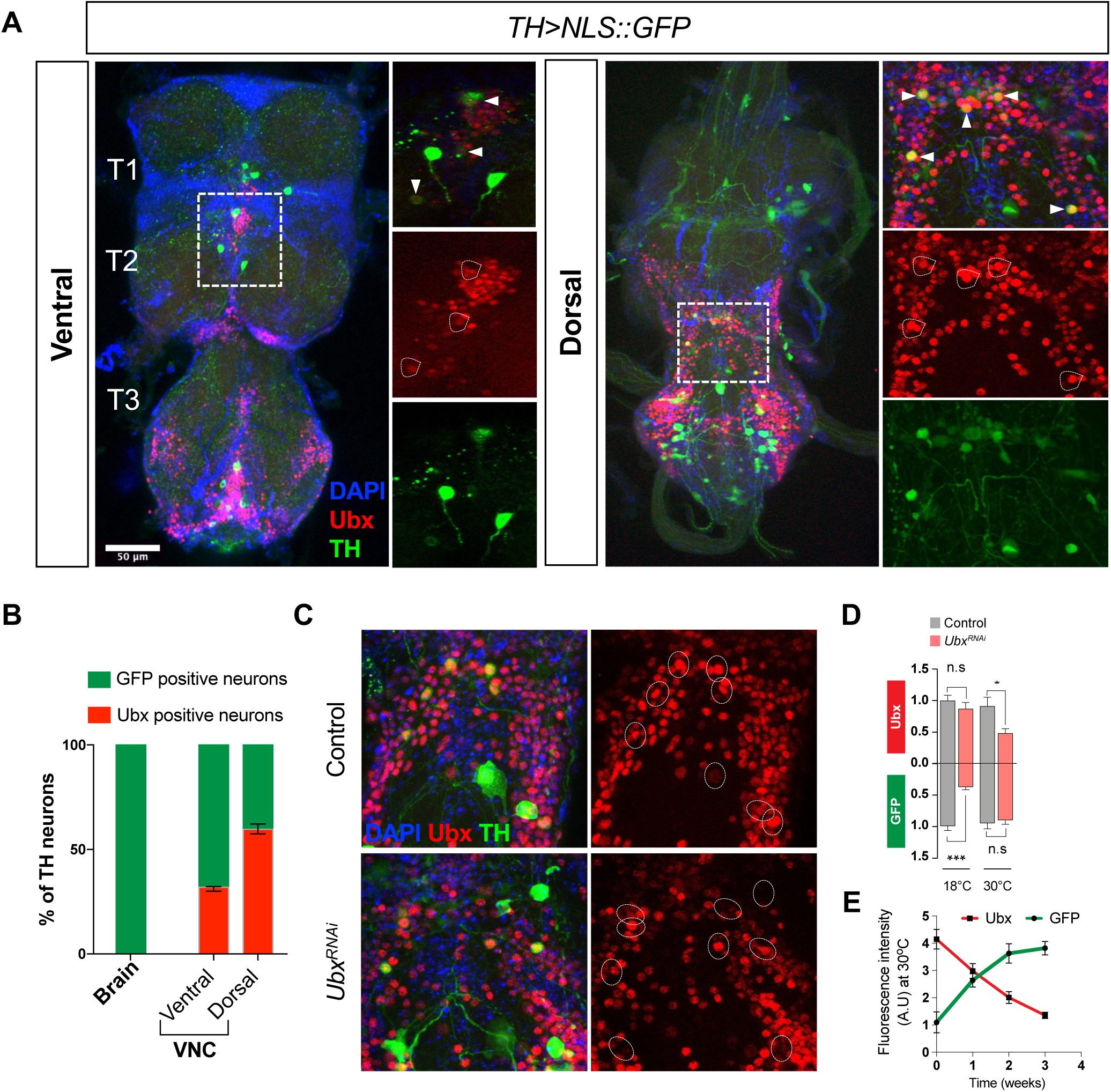
Ubx is expressed in adult VNC Dopaminergic (TH) neurons. (**A**) Confocal images of the VNC of adult *Drosophila* showing expression of Ubx expression in TH neurons labeled by GFP driven by *TH-Gal4*. (**B**) Ubx is expressed in ∼ 30% (5±0.9) and 60% (17±0.5) of TH neurons located respectively in ventral and dorsal sides of the VNC (n=8-10). (**C-E**) Conditional reduction of Ubx expression by *Ubx^RNAi^* in TH neurons. (**C**) Confocal imaging of a region of the adult VNC with high Ubx protein expression, showing the effects of *Ubx^RNAi^* treatment at 30°C on Ubx expression. Genotypes: control (*TH-Gal4; UAS-NLS::GFP, UAS-FLP*); *Ubx^RNAi^* (*TH-Gal4; UAS-NLS::GFP,Tub-Gal80ts; UAS-Ubx^RNAi^*). Note the apparent decrease in red signal within TH neurons (White circles) (Flies were 8-10 days old, n=5-6). (**D**) Quantification of Ubx expression in normal and *Ubx^RNAi^* conditions at permissive (18°C) and non-permissive (30°C) temperatures for Gal80ts repression. Note that, at low temperature, Ubx levels are not different between control and *Ubx^RNAi^* lines; in contrast, GFP expression is highly reduced due to Gal80ts repression. At high temperatures, Ubx expression is significantly downregulated. (**E**) Kinetics of Ubx and GFP expression in adult flies expressing *Ubx^RNAi^* in TH neurons (*TH-Gal4; UAS-NLS::GFP,Tub-Gal80ts; UAS-Ubx^RNAi^*) in non-permissive conditions (30°C) over time (1-3 weeks). Note that as time elapses the reduction in Ubx expression becomes more pronounced, and, in contrast, expression of GFP becomes stronger (E). Error bars represent SEM. Significant values in all figures based on Mann-Whitney U test: ^∗^p < 0.05.

We then examined the relation between the activity of the dopaminergic system and flight control using optogenetics (*19*). For this, we firstly expressed CsChrimson(*19*) in the TH-domain and monitored effects on flight, observing that optogenetic activation of TH neurons (N.B. in the absence of an air-puff) is sufficient to initiate flight (Fig. 3A and Movie 4). Secondly, we optogenetically inhibited TH neurons using GtACR(*20*) and observed that this treatment is sufficient to significantly reduce wing flapping (Fig. 3B and Movie 5). These results demonstrate that modulation of neural activity of TH neurons affects flight. Similar flight suppression effects were also observed upon optogenetic treatment of flies expressing GtACR within the Ubx domain (Fig. 3C and Movie 6) (NB: expression of CsChrimson within the Ubx domain leads to lethality). These observations demonstrate a functional link between the activity of the dopaminergic system and flight maintenance.

**Fig. 3.**
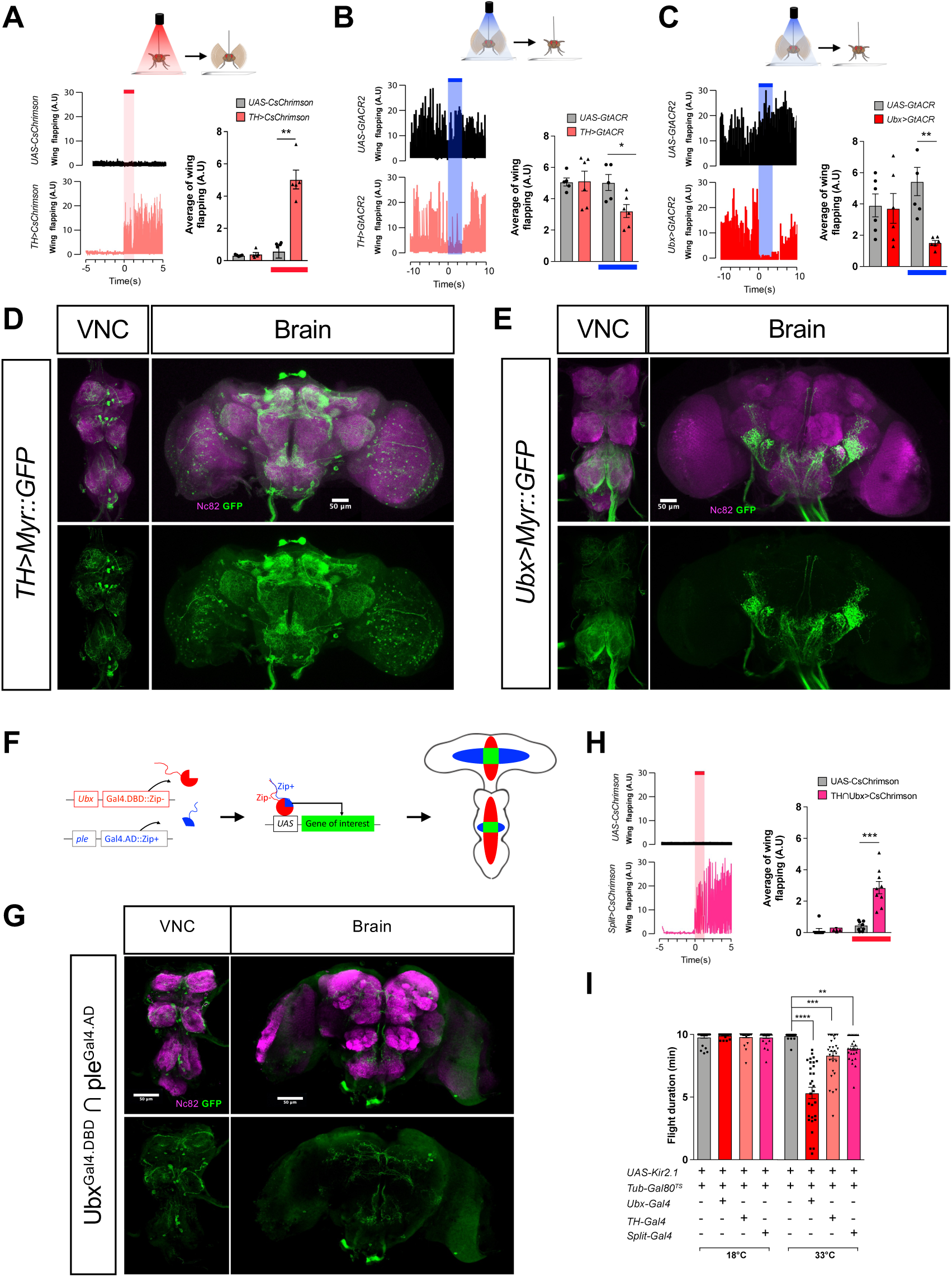
Modulation of activity in dopaminergic neurons expressing Ubx affects flight. (**A**) Optogenetic activation (red shade) of TH neurons expressing channelrhodopsin CsChrimson (*TH>CsChrimson*) induces spontaneous flight (Top: cartoon representation of experimental results) (**B-C**). Optogenetic inhibition (blue shade) of TH neurons (*TH>GtACR2*) (B) and Ubx+ cells (*Ubx>GtACR2*) (C) by expressing GtACR reduce flight. In each figure, left panels represent wing flapping frequency, and right panels depict a histogram of average frequencies. Control flies are *UAS-CsChrimson* and *UAS-GtACR* (n=6). (**D**-**E**) Confocal images showing TH (D) and Ubx cells (E) in VNC and brain. (**F**) Schematic illustration of the Split-Gal4 system based on the complementation between the two functional domains of Gal4, the DNA-binding (DBD) and transcription-activation (AD) domains. Each domain is fused to a heterodimerising leucine zipper (Zip+ or Zip-) that promotes the fusion of the two domains when expressed in the same cell reconstituting transcriptional activity. (**G**) Confocal images of the fly VNC (ventral side) and brain (anterior side) showing the UAS-Myr::GFP expression pattern driven by *Ubx^Gal4.DBD^ ∩ ple^Gal4.AD^* defines a subset of TH neurons that express Ubx. (**H**) Flight patterns of flies before and after optogenetic activation of Ubx^+^ TH cells, (n=8-9). (**I**) Flies with Ubx positive neurons inhibited by expression of the potassium channel encoded by the *Homo sapiens* KCNJ2 gene (Kir2.1) show reduced flight duration; slight reduction is also observed when Kir2.1 is expressed in the TH-Gal4 domain, and, notably, in the Ubx*∩*TH intersectional domain (split-Gal4). See also Movie 4 (related to Fig.3A), Movie 5 (related to Fig.3B), Movie 6 (related to Fig.3C), and Movie 7 (related to Fig.3H). Error bars represent SEM. Significant values in all figures based on Mann-Whitney U test or one-way ANOVA with the post hoc Tukey-Kramer test: ^∗^p < 0.05, ^∗∗^p < 0.01, and ^∗∗∗^p < 0.001.

To gain further insight on the properties of Ubx^+^ dopaminergic neurons in connection to flight control, we developed a split-Gal4 approach (*21*). For this, we expressed complementary forms of the Gal4 transcriptional activator from two distinct promoters: *Ubx* (Gal4.DBD::Zip-) and *pale* (*ple*, the gene that encodes TH) (Gal4.AD::Zip^+^) in order to reconstitute functional Gal4 protein only at the intersection between the *Ubx* and *TH* transcriptional domains (Fig. 3F-G and Figure S3A); this design enabled us to develop a range of experiments to test the functional roles of *Ubx^+^* TH neurons in relation to flight control. First, optogenetic activation of TH∩Ubx (split-Gal4, “split”) neurons is sufficient to trigger flight in the absence of an air-puff (Fig. 3H and Movie 7). Second, FlpOut-induced expression of red activatable channelrhodopsin (ReaChR) only within the Ubx^+^ TH domain (via *TH>LexA;LexAOP-FLP* and under *Ubx-Gal4* induction) (Fig. S3B), triggers flight under red-light illumination. This further demonstrates that the activity of Ubx^+^ dopaminergic neurons is linked to flight control (Fig. S3C,D). Third, thermogenetic inhibition of neural activity mediated through expression of the inward-rectifier potassium ion channel Kir2.1/KCNJ2(*22*) leads to a significant reduction of flight duration when applied to TH∩Ubx (“split-Gal4”) neurons (Fig. 3I). These experiments indicate that activity levels of Ubx^+^ TH neurons are directly related to flight regulation: inhibition of these neurons halts flight, while their stimulation triggers flight. Building on our neuronal activity manipulations, and given that *Ubx* expression (*Ubx^RNAi^*) within TH-neurons does not induce a visible change in the number of TH neurons (Fig. S4A,B), we sought to determine the natural patterns of activity of TH neurons in relation to normal flight, and how these were affected when *Ubx* expression was reduced, exploring the possibility that changes in Hox expression in the adult, might affect neural physiology. For this we made use of the CaLexA system (calcium-dependent nuclear import of LexA) (Fig. 4A)(*23*), which acts as a recorder device of neural activity in the tissue/cellular context of choice, and, thus, provides a platform to correlate neuronal activity patterns with specific behaviours (*24*). In a first series of experiments, we drove the CaLexA reporter from the *TH-Gal4* driver and observed that after 30-min of constant flight there is a significant increase of CaLexA signal in the dopaminergic system when compared to a resting condition (Fig. 4B-E). Indeed, CaLexA signal intensity within the TH domain, is positively correlated with flight duration (Fig. 4F) indicating that the dopaminergic system is active during flight. In a second series of experiments, we reduced *Ubx* expression (by means of *Ubx^RNAi^*) within TH neurons (although these experiments were conducted across the whole TH domain, expression of *Ubx* could have only been reduced in those cells that normally express the gene), and observed a significant decrease of neural activity in both, the second (T2) and third (T3) thoracic segments (Fig. 4G-I). These data strongly support the notion that a reduction of *Ubx* expression in dopaminergic neurons leads to a decrease in neural activity within this domain. Interestingly, although we can indeed detect dopaminergic neurons in the T1 segment, as well as in the brain (Fig. S4C), we do not observe any CaLexA signal after flight in these regions. Further results indicate that flight-related dopaminergic activity takes place primarily within the VNC in segments T2-T3 (Fig. 4J), the thoracic regions where neural expression of *Ubx* is prominent (*3, 10*). Expression of the CaLexA reporter within the intersectional domain TH∩Ubx reveals that even within this restricted aspect of the dopaminergic system, there are significant differences in CaLexA signal detected after flight, when compared with a resting control (Fig. 4K-M). Furthermore, expression of the bacterial voltage-gated sodium channel NaChBac(*25, 26*) within dopaminergic neurons expressing reduced levels of *Ubx*, leads to a rescue of the flight phenotype observed after TH-driven *Ubx^RNAi^*, as assessed by two independent behavioural tests: tethered flight (Fig. 4N and Fig. S4D) and forced flight (Fig. 4O); this latter observation strongly suggests that a reduction of Ubx affects the physiological properties of the neuron, rather than its morphology. Altogether, the data above indicate that normal *Ubx* expression within the thoracic dopaminergic system is required for flight-related neural activity in thoracic TH neurons.

**Fig. 4.**
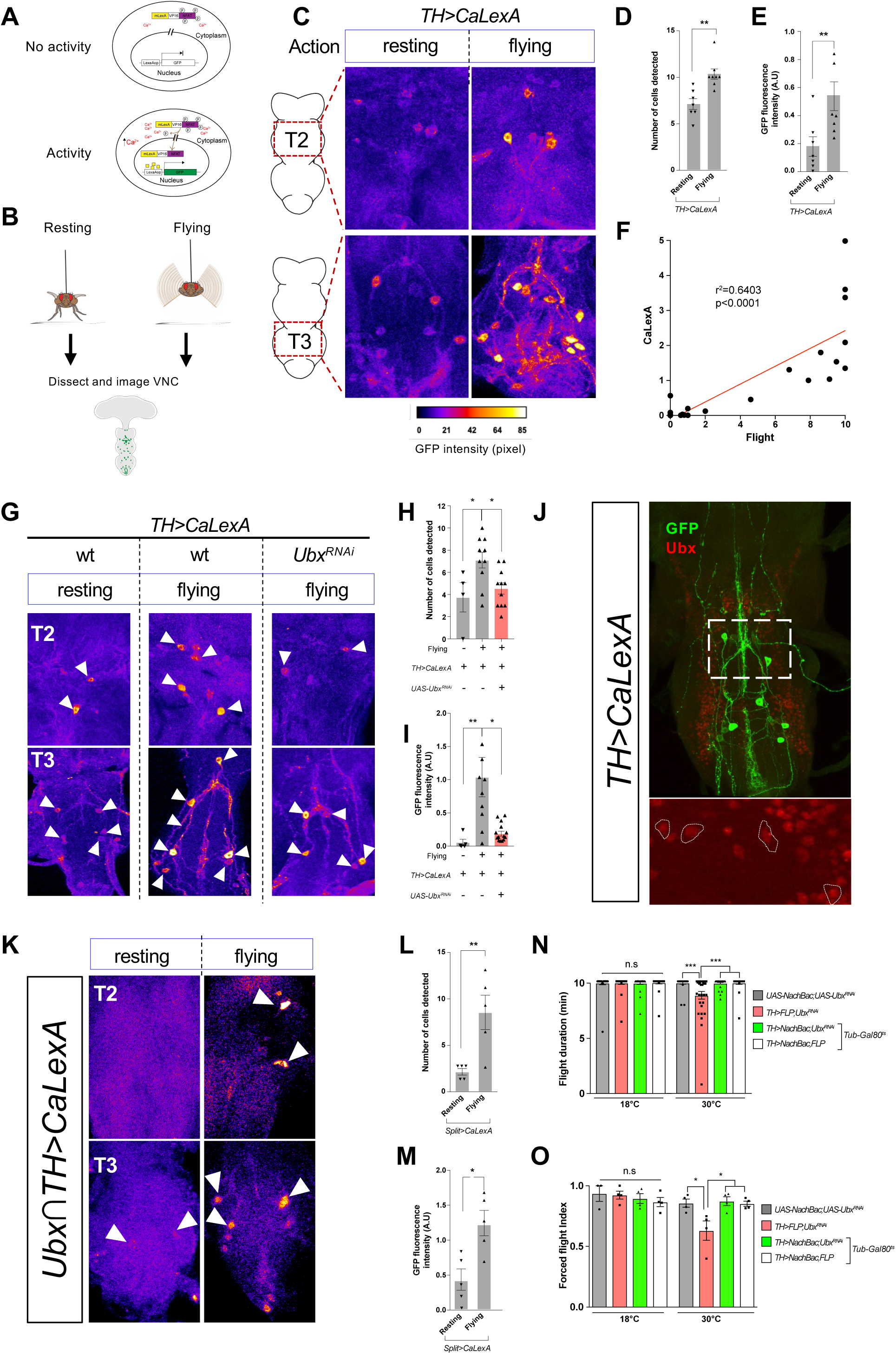
Levels of neural activity of *Ubx^+^* TH neurons during flight. (**A**) Schematic illustration of the CaLexA system, which records neuronal activity based on Ca^2+^-NFAT interaction. Cartoon representation of the CaLexA system in the absence (top panel) and presence of neural activity (bottom panel); in the latter, Ca^2+^ accumulation dephosphorylates NFAT, triggering the transport of the transcription factor mLexA-VP16-NFAT into the nucleus, where the chimeric transcription factor LexA binds to the LexA operator (LexAop), and subsequently induces expression of the GFP reporter gene. (**B**) Experimental protocol. Resting and flying Drosophila were collected for dissection and their VNCs were immunostained and imaged. (**C**) Representative images of TH neurons in the VNCs of resting and flying wild-type *Drosophila* (TH>CaLexA), immunolabelled with GFP (warmer color). Note that resting flies show low GFP intensity/CaLexA signals while flying flies show very high signal intensity. (**D-E**) Quantification of GFP positive TH neurons (D) and GFP intensity (E) of the VNC thoracic segments 2 (T2) and 3 (T3) in *TH>CaLexA* flies in different conditions (resting: n=7; flying: n=8). (**F**) Increase in GFP intensity is correlated with flight (n = 21). (**G**) Representative images of TH neurons in the VNCs of wild-type (wt) *Drosophila* (*TH>CaLexA*) and TH neurons expressing *Ubx^RNAi^* (*TH>CaLexA; Ubx^RNAi^*). (**H-I**) Quantification of GFP positive TH neurons (H) and GFP intensity (I) of VNC in segments T2 and T3 in *TH>CaLexA* and *TH>CaLexA; Ubx^RNAi^* (n>5). (**J**) GFP positive TH neurons are also Ubx positive. (**K**) Representative images of the Ubx*∩*TH intersectional domain in the VNC of resting and flying *Drosophila* (*Ubx∩TH >CaLexA*). (**L-M**) Quantification of GFP positive cells (L) and GFP intensity (M) of VNC in T2 and T3 segments in *Ubx∩TH >CaLexA* (n=5). (**N**-**O**) Increase in TH neurons activity generated by conditionally expressing the voltage-gated bacterial Na^+^ channel (NaChBac) rescues the flight deficit resulting from Ubx downregulation. Averages of flight duration (N) and forced flight index (O) of flies after heat shock (at 30°C). Error bars in figures represent SEM. Significant values in all figures based on Mann-Whitney U test or one-way ANOVA with the post hoc Tukey-Kramer test: ^∗^p < 0.05, ^∗∗^p < 0.01, ^∗∗∗^p < 0.001.

To probe the relation between the dopaminergic system and the flight motor system underlying normal flight, we first considered the (unlikely) possibility that thoracic TH neurons innervate flight muscle directly. Expression of membrane-bound GFP in all TH neurons shows that they do not produce any direct contacts with the flight motor (Fig. 5A) indicating that dopaminergic neurons must be communicating with the flight muscle system indirectly. To test this model, we reduced expression of the dopamine receptors (Fig. 5B) exclusively in those motor neurons that innervate the dorsal longitudinal muscle field in the thorax (directly controlling wing flapping) by means of drivers targeting flight muscle motoneurons (*15*) (Fig. 5C,D and Fig. S5A-C). To achieve this we expressed *UAS-Damb^RNAi^* – known to reduce expression of the gene encoding Dop1R2/DAMB(*27, 28*), one of the dopamine receptors – under control of *Dlm-Gal4* (*VT021842.GAL4.attp2*), which is active in dorsal longitudinal muscles (DLMs) (Fig. 5C,D). This perturbation resulted in a significant reduction in flight ability (Fig. 5C,D) providing support to the notion that thoracic TH neurons are pre-synaptic to flight command motor neurons. In contrast, reduction of expression of the dopamine receptor in other motor neurons involved in flight control, as driven by the *tp1, tp2-Gal4, tt-Gal4 and tp2-Gal4* lines does not significantly alter flight duration as assessed in tethered flight experiments (Fig. 5C,D). These observations suggest that TH neurons control the activity patterns of Dlm motor neurons (VT021842 motor neurons), and that these motor neurons are directly related to flight control. To confirm this, we optogenetically activated Dlm neurons using CsChrimson and observed that this treatment indeed activated wing flapping (Fig. 5E, and Movie 8). In addition, thermogenetic inhibition of neural activity of Dlm neurons using Kir2.1/KCNJ2 (Fig. 5F and Fig. S5D), or blocking synaptic transmission of these neurons by expressing a dominant-negative form of the dynamin GTPase, *shibire^ts^* (Fig. S5E,F) lead to a reduction in flight duration indicating that the Dlm motor neurons are directly involved in flight maintenance. These data are consistent with previous observations on the links between these motor neurons with the frequency and the number of wing beat cycles during courtship, and flight ability (tested by voluntary take-off and in a drop assay) (*15*). Furthermore, membrane-tagged GFP expressed in Dlm motor neurons shows that these motor neurons innervate the longitudinal muscles in the fly thorax, providing a direct neuronal link between the dopaminergic system and flight muscles controlling flight (Fig. 5G)(*29*). To confirm structural connectivity between the TH neurons and DLM motor neurons we used the GRASP system (*30, 31*) (Fig. 5H) expressing complementary forms of GFP in both putative pre-synaptic (TH-Gal4) and post-synaptic (DLM-Gal4) elements, and observed reconstitution of functional GFP signal in both T2 and T3 segments (Fig. 5I) providing strong structural evidence of a direct link between the dopaminergic system, and the Dlm motor neurons commanding flight. Furthermore, optogenetic stimulation of TH neurons leads to activation of Dlm motor neurons (Fig. 5J-K and Movie 9) demonstrating a functional, physiological coupling between these neurons. Altogether, this dataset provides a cellular framework that explains how reductions in the activity of the dopaminergic system relate to alterations in flight.

**Fig. 5.**
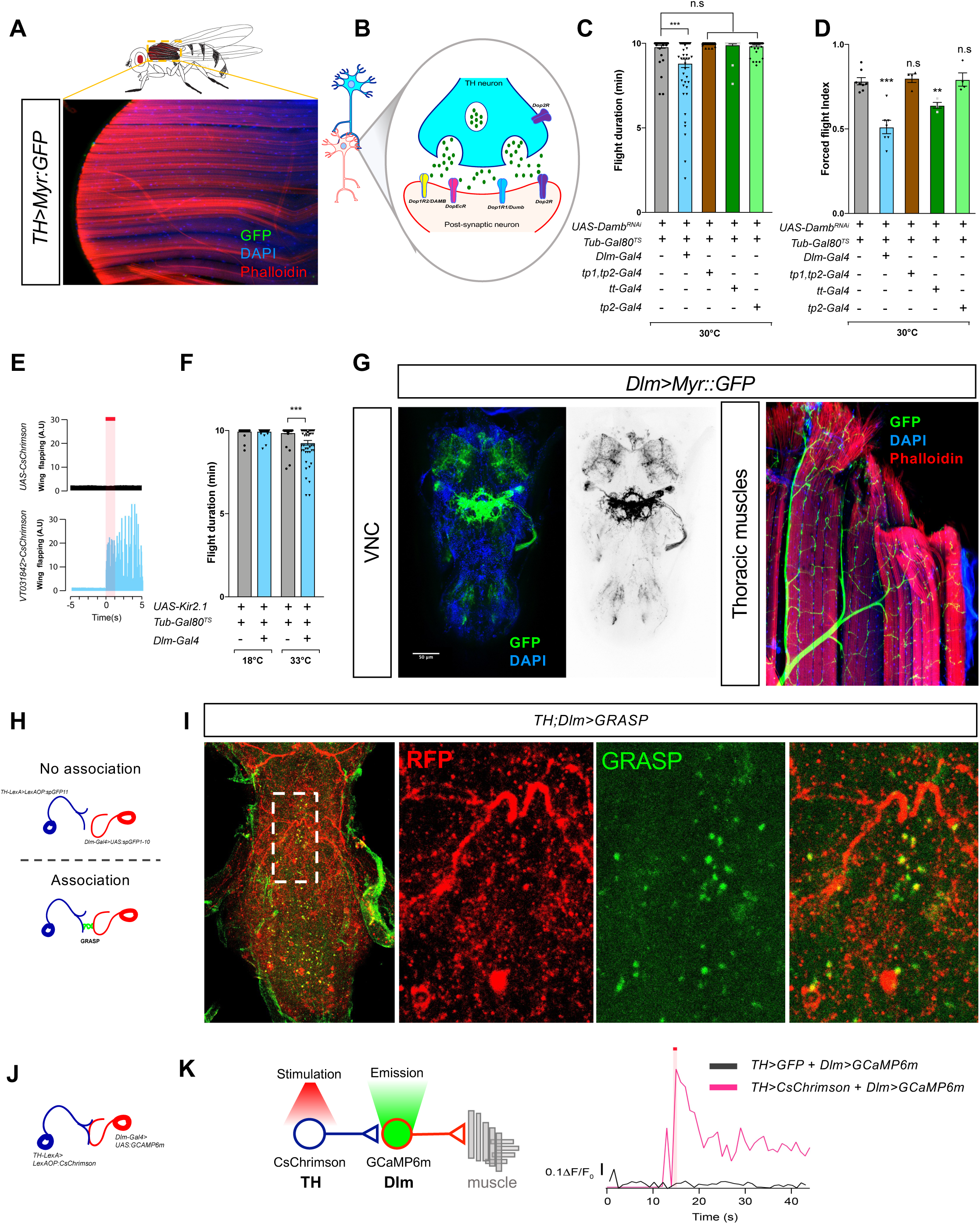
H neurons modulate flight through direct contact to flight motoneurons. **(A)** Confocal image of thoracic muscles of fly expressing myristoylated GFP in dopaminergic neurons (*TH>Myr:GFP*). No GFP signal can be observed. (**B)** Schematics of a dopaminergic synapse and receptors. (**C-D**) Average flight duration (C) and force flight (D) at 30°C of flies with downregulated expression of dopamine receptors (*DAMB^RNAi^*) in specific flight motor neurons. (**E**) Red light-evoked (red shade) activation of dorsal longitudinal flight muscle (Dlm) motoneuron expressing CsChrimson (*Dlm>CsChrimson*) induces spontaneous flight (n≥ 40). See also Movie 8. (**F**) Flies with Dlm neurons inhibited by expression of Kir2.1 show reduced flight duration. (**G**) Representative confocal images showing the projections of Dlm motoneurons within the VNC (left panel) and longitudinal flight muscles (right panel). (**H**) Cartoon illustrating the principle underlying GFP Reconstitution Across Synaptic Partners (GRASP). GFP is reconstituted when two complementary segments of GFP associate on the extracellular surfaces of adjacent neurons. (**I**) Confocal image of GRASP reconstitution in the VNC of *Drosophila TH;Dlm>GRASP* (*TH-LexA>LexAOP:RFP,LexAOP:spGFP11 & Dlm-Gal4>UAS:spGFP1-10*). GRASP fluorescence reveals structural links between TH and Dlm neurons in the VNC. (**J**) Cartoon illustrating the experimental approach to determine a functional connection between TH and Dlm neurons. (**K**) Red light-evoked (red shade) activation of TH neuron expressing CsChrimson (*TH>CsChrimson*) induces spontaneous activity of Dlm in *Drosophila* TH;Dlm>CsChrimson; GCAMP6 (*TH-LexA,LexAOP:CsChrimson;Dlm-Gal4,UAS:GCAMP6m*). See also Movie 9. Error bars represent SEM. Significant values in all figures based on Mann-Whitney U test or one-way ANOVA with the post hoc Tukey-Kramer test: ^∗^p < 0.05, ^∗∗∗^p < 0.001.

To advance the understanding of the mechanisms that link a reduction in *Ubx* expression with the observed physiological change in TH neurons with impact on flight performance (Fig. 1, and Fig. 4), we conducted an RNA-sequencing experiment aimed at determining the transcriptome of TH neurons from the adult VNC. For this, we used FACS to isolate populations of TH neurons from the VNC expressing normal or downregulated *Ubx* (*TH>Ubx^RNAi^*), extracted RNA, and compared the resulting transcriptomes using RNA-seq (Fig. 6A). Using edgeR analysis(*32*) we identify 233 differentially expressed genes (DEGs) (out of 5709 total genes detected) in *TH>Ubx^RNAi^* neurons relative to wt neurons (p-value <0.01; Table S1) (Fig. 6B). Using the DAVID platform (Database for Annotation, Visualization and Integrated Discovery(*33*)) we established the functional biological properties of the 233 DEGs. Looking for candidate genes that might perform *Ubx*-dependent physiological roles in TH neurons, we noted that amongst the top-ten DEGs (Fold change (FC) >5) were four previously uncharacterised ion transport genes, with predicted symporter activity related to sodium, calcium, or potassium ion transport (Fig. 6C) (i.e. Gene Ontology (GO) terms enriched, Holm-Bonferroni test, p-value<0.05). These genes, which are substantially downregulated in response to a reduction of *Ubx* expression (≥10 FC, p<0.004; Fig 6D), include: *CG1090*, *CG5687*, *CG9657* and *CG6723* (Fig. 6D). Gene tree (Fig. 6E), protein alignment analysis (Fig. S6 and S7), and protein structural predictions (Fig. 6F and Fig. S6, 7) using EMBL-EBI Clustal Omega, Jalview program(*34*), and JPred secondary structure prediction programs (*35*) for these symporter genes reveals that they belong to two independent lineages within the Solute Carrier gene (SLC) family: SCL5 and SCL24 (*36–39*). Although these genes have not been previously characterised in flies, the SLC24 genes have been shown to encode a diverse group of Na^+^/Ca^2+^-K^+^ exchangers (NCKX)(*37*) in other species, and have been previously shown to play roles in nutrient sensing and sleep control in insects and mammals(*36, 37, 39, 40*), and their dysfunction is correlated with neurological disease (*36, 41, 42*). Furthermore, expression data from the FlyAtlas expression database (*43*), reveals that these four symporter genes are primarily expressed in the *Drosophila* CNS, including the VNC (Table S2). Put together, these features suggest the hypothesis that SCL genes might play functional roles in the fly nervous system and might be some of the mediators through which Ubx exerts its roles on flight. To test whether normal expression of these symporter genes in TH neurons is required for flight maintenance, we examined flight behaviour (tethered flight and forced flight) following TH-specific RNAi-mediated downregulation of these genes. Our results show that TH-specific expression of RNAi constructs against *CG1090* and *CG6723* lead to an impairment of flight maintenance and ability (Fig. G,H and Fig. S8) indicating that normal expression of these symporter genes in TH neurons is necessary for normal flight. Altogether, these experiments suggest a molecular framework that links *Ubx* gene-regulatory roles in TH neurons to flight control.

**Fig. 6.**
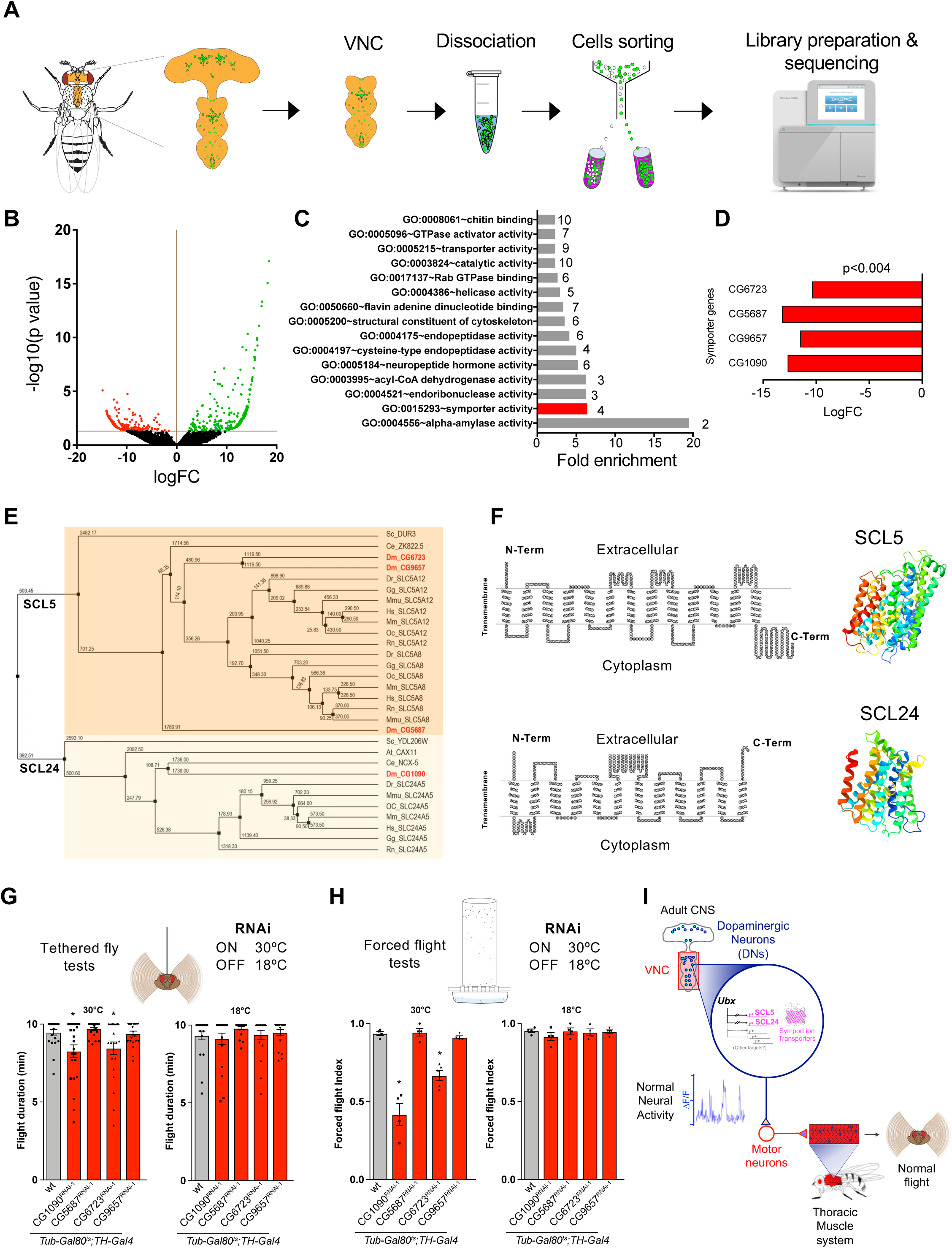
Solute-carrier gene (SLC) symporter genes are differentially expressed in response to Ubx downregulation in TH neurons. (**A**) Experimental design of our neuron-specific transcriptomic experiment. (**B**) Volcano plot depicting differentially expressed genes (DEGs). Red and green dots represent down- and upregulated genes, respectively. (**C**) Gene Ontology (GO) enrichment analysis. GO molecular functions of DEGs show top four genes (red bar) predicted to have symporter activity (involvement in calcium, potassium, sodium and/or solute co-transport). (**D**) The four differentially expressed symporter genes are strongly downregulated. None of these genes has been previously characterised in flies. (**E**) Dendrogram of the differentially expressed symporter genes detected in *Drosophila* showing their relation to other gene families across the animal kingdom. The analysis suggests that the differentially expressed symporter genes belong to the human SLC5 and SLC24 protein families. Abbreviations are: *Saccharomyces cerevisiae* (Sc), *Arabidopsis_thaliana* (At), *Danio rerio* (Dr), *Gallus gallus* (Gg), *Homo sapiens* (Hs), *Macaca mulatta* (Mm), *Oryctolagus cuniculus* (Oc), *Rattus norvegicus* (Rn), *Mus musculus* (Mmu) *Drosophila melanogaster* (Dm), *Caenorhabditis elegans* (Ce) (values near/above internodes correspond average of branch lengths from T-Coffee analyses). (**F**) Structural predictions for the symporter proteins encoded by differentially expressed symporter genes based on human SLC5A8 and SLC24A5. Left panel: SACS MEMSAT2 graphs predicting 13 and 12 transmembrane domains for symporter proteins SLC5 and SLC24, respectively. Right panel: predicted 3D structures of SLC5 and SLC24 proteins. (**G**-**H**) Conditional downregulation of symporter genes CG1090 and CG6723 in TH neurons affect flight maintenance in tethered flies (G) and ability in forced flight experiments (H). (**I**) Proposed cellular and molecular model of Ubx-dependent control of flight behaviour in *Drosophila*. Error bars in figures represent SEM. Significant values in all figures based on Mann-Whitney U test or one-way ANOVA with the post hoc Tukey-Kramer test: ^∗^p < 0.05.

Our work reveals that normal *Ubx* expression in the adult nervous system is required for normal flight, revealing a novel post-developmental role of the *Hox* genes in adult behaviour. These observations open a new avenue to investigate the molecular programmes that maintain normal adult neural physiology, and based on the broad evolutionary conservation of the *Hox* system across distinct animal taxa, suggest that the *Hox* genes may play neurophysiological roles in the adult forms of other species, including humans.

## ACKNOWLEDGEMENTS

We wish to thank Aishwarya Padmanabhan and Jonathan Menzies for their contributions to neuronal dissections, and all members of the Alonso Lab for helpful discussions and feedback on this work. We also thank Takeshi Yoshimatsu and Andre Chagas for their assistance with Igor software. We are very grateful to Scott Waddell, Serge Birman, the Vienna Drosophila Resource Center, and the Bloomington Stock Centre for fly stocks. This research was funded by a Wellcome Trust Investigator Award (098410/Z/12/Z) made to C.R.A., and a Medical Research Council Project Grant (Ref: MR/S011609/1) given to C.R.A..

## MATERIALS AND METHODS

### Fly rearing and stocks

Drosophila cultures were reared at 25 °C in standard culture tubes or bottles, and on corn agar media supplemented with water, cornmeal, molasses, yeast, nipagin, and propionic acid. The following Gal4 drivers from Bloomington *Drosophila* Stock Center (BDSC) were used: Pan-neuronal Gal4 (Elav-Gal4), cholinergic (Cha-Gal4)(BDSC, 60317), Glutamatergic (VGlut-Gal4) (BDSC, 26160) and Ubx-Gal4 from BDSC Gift from Ernesto Sánchez-Herrero(*44*). The dopaminergic GAL4 (TH-GAL4) were generously provided by Serge Birman (CNRS, ESPCI Paris Tech, France)(*31*). In addition, various GAL4 drivers for motoneurons innervating flight muscles(*15*) were used, VT021842.GAL4 (VDRC, 203853), VT033051.GAL4 (VDRC, 200860), VT040232.p65ADZp.attp40 (BDSC, 71651), VT029310.ZpGAL4DBD (DBSC, 74584), VT045969.p65ADZp.attp40 (BDSC, 73438), and VT042475.ZpGAL4DBD.attp2 (BDSC, 72758). We combined UbxDBD/R32D08-Gal4.DBD(BDSC, 68670), ple-p65.Gal4.AD (BDSC, 70861) to make split-Gal4 lines. We used UAS strains of dopamine receptor RNAis : DR2/Dop1R2/DAMB (VDRC, 105324, BDSC, 26018), DR1/Dumb/Dop1R1 (VDRC, 107058), D2R/Dop2R (VDRC, 11470, 11471), and DopEcR (VDRC, 103494), and Ubx-RNAi (31913 and 34993, referred respectively to as Ubx-RNAi and UbxRNAi-2 in the article), as well as mutant Ubx (REF) a Tub-GAL80^ts^(BDSC, 7018 and 7108). *UAS-n-syb::spGFP_1-10_*, *LexAop-CD4::spGFP_11_*/*CyO* and *LexAop-n-syb::spGFP_1-10_*, *UAS-CD4::spGFP_11_* (GRASP)(BDSC, 58755 and 58754)(*45*). We obtained Mef2-LexA from Scott Waddell (Centre for Neural Circuits and Behaviour, University of Oxford, Oxford)(*46*). TH-LexA from Ronald L. Davis provided by Serge Birman (CNRS, ESPCI Paris Tech, France)(*47*), UAS-GtACR from Adam Claridge-Chang (Program in Neuroscience and Behavioral Disorders, Duke-NUS Medical School, Singapore)(*48*), 8XLexAop2-FLPL (BDSC, 55819), UAS-CsChrimson (BDSC, 55819 and 55135), UAS-NaChBac (BDSC, 9469), UAS>stop>ReaChR(BDSC, 53743)(*49*), UAS-Kir2.1 (BDSC, 6595), UAS-Myr::GFP (BDSC, 31913), Kir2.1 (BDSC, 6595), UAS-UbxFLP (42719, 42720) and UAS-Shibire^ts^ (BDSC, 44222) (*50*). UAS-Nls:GFP (BDSC, 4776); lexAop-CD8-GFP-2A-CD8-GFP;UAS-mLexA-VP16-NFAT,LexAop-rCD2-GFP (CaLexA) (BDSC, 66542), UAS-GAMP6 (BDSC, 42748)(*51*). w1118 was used as control strain.

### Flight assays

For tests conducted on tethered *Drosophila*, flies were anaesthetised on ice for 15 min and tethered to the tip of a thin metal wire (Insect pins 0.1mm, Agar Scientific) with UV-activated glue (BONDIC). After 20 min of acclimatisation at room temperature, a brief air-puff was delivered to the animal, and subsequently videos were recording during 10 min via a Basler ACE acA1300-200uc Colour USB 3.0 Camera and TECHSPEC® 6mm UC Series Fixed Focal Length Lens, with a resolution of 1,280 x 1,024 pixels at 50 Hz to measure flight duration for each genotype. For forced flight assays, we proceeded as previously described (Fig. 1F) (*52*). We tapped groups of (n ≥12) flies (males/females) through a funnel in a transparent plastic cylinder (90 cm high, 13.5 cm inner diameter) into which we introduced a folded polyacrylamide sheet (90 cm high) coated with Tangle-Trap (The Tanglefoot Company), an adhesive designed to capture live insects. A bucket with water was placed underneath the cylinder. Five minutes after the flies were released the plastic was removed from the cylinder, unfolded coated faceup on a flat white surface, and its top view captured by a digital camera. Lastly, FIJI image J software was used to automate the scoring of flight performance. The polyacrylamide sheet was divided in 3 zones: top> 80 cm, bottom < 2 cm, and bottom <MEDIUM>top, and forced flight index was scored according to the following formula: (n_top_-n_bot_)/2n_tot_, where n_tot_ is the total number of flies, and n_top_ and n_bot_ the number of flies at the top and at the bottom, respectively. The number of flies landing in the bucket were included in the ‘bottom’ group. For conditional expression of UbxRNAi, Kir2.1 or NachBAC transgene in the *TubGAL80^t^*^s^ background, animals were shifted from permissive (18°C) to non-permissive (30°C) temperature upon eclosion and maintained at 30°C until the experimental day. Experiments were performed at room temperature (approximately 25°C). For voluntary flight assays, 4-6 flies were introduced one by one into 5 cm diameter circular arena with sloped edges (11°) covered with glass that was pre-coated with Sigmacote (SIGMA-ALDRICH). This arena triggers an escape reflex to the point that lifting the glass cover allows flies to immediately take-off from the open container. We next, lifted the lid of the arena and after 2 min take-off was noted as successful for the flies that did not land within a radius of 7 cm.

Optogenetic experiments were conducted by adapting Flypi device(*53*). For neuronal activation (CsChrimson, Pwr590) and inhibition (GtACR, Pwr_470_) a Neopixel twelve light-emitting diodes ring was positioned face-down around the infrared-capable camera objective about 3 cm above the tethered flies. Flies were recorded with lights off (in dark) and the response of the animal (males and females) were analysed during the following light on period. We exposed flies to approximately 4.9W/cm^2^ for stimuli between 500-1000 ms using a custom-written Graphical User Interface (*53*). For all optogenetic activation experiments, adult flies upon eclosion were kept for 7-8 days before the experiment on food containing 0.5 mM all-trans retinal (Sigma).

### Climbing assays

Startle-induced negative geotaxis (SING) test was used as climbing assay, and was monitored as previously described (Issa et al., 2018). A group of 10-20 flies were introduced in a vertical glass column (15cm long, 1.5cm inner diameter) with a flat bottom end. After approximately 25 min of acclimatisation flies were tested individually by gently tapping them down (startle), and scoring the number of flies having reached the top of the column (above 12 cm) and remaining at the bottom end (below 2 cm) after 25s. Each group was assayed 3 times at 10 min intervals. The climbing performance for each column was calculated as follows: (n_tot_ + n_top_ -n_bot_)/2n_tot_, where n_tot_ is the total number of flies, and n_top_ and n_bot_ the number of flies at the top and at the bottom, respectively. Results are the mean and SEM of the scores obtained with the 5 groups of flies per genotype.

### Immunohistochemistry

Adult ventral nerve cords (VNC) were dissected in 1X PBS. Tissues were then fixed for 1h in 4% formaldehyde in 1X PBS at room temperature. After fixation, brains and VNCs were washed 3 times (30 min per washing) in PBS with 0.3% Triton X-100 (PBTx) and incubated at 4°C overnight with primary antibodies. The following primary antibodies were used: mouse monoclonal anti-Ubx (FP3.38(*3*) 1:500) from the Developmental Studies Hybridoma Bank), rabbit anti-TH (Novusbio), mouse anti-nc82, and chicken anti-GFP (Abacam Probes, 1:3000). The secondary antibodies were anti-mouse Alexa Fluor 555 (Invitrogen Molecular Probes, 1:1000), anti-rabbit Alexa Fluor 647 (Invitrogen Molecular Probes, 1:1000) and anti-chicken Alexa Fluor 488 (Invitrogen Molecular Probes, 1:1000). Images were acquired with a Leica SP8 confocal microscope, processed, and analysed using FIJI ImageJ.

### Calcium activity recording in thoracic muscles and CaLexA assays

Calcium activity recording in thoracic muscles was conducted on tethered fly, and calcium signalling recorded using epifluorescence microscopy Leica DM6000. Next, FIJI software, and subsequently Igor program were used to quantify calcium signal and generate traces, respectively. The CaLexA (Calcium-dependent nuclear import of LexA) technique (*23*) was employed to measure calcium activity in the dopamine and Ubx neurons. Briefly, CaLexA expressing flies were raised at 18°C. After eclosion temperature was shifted to 30°C, and males and females were separated and put into fresh vials. To restrict fly movement, a cotton plug was placed close to the surface of food and vials were covered by aluminium foil. The day before the experiment animals were moved to 25°C for acclimatisation. The day of the experiment animals were cold-anaesthetised on ice for 15 min prior to tethering. A group of tethered flies were allowed to hold on the ground with their legs to prevent spontaneous flight (resting condition) in contrast to others, which were kept on suspension (‘flying’ condition). A single blown air-puff stimulus was delivered to the animals to trigger flight (a single brief air-puff stimulus is sufficient to trigger flight in tethered *Drosophila*). After 30min of flight, or cessation of the flight, animal VNCs were dissected and processed for immunohistochemistry as described above. Samples were visualized with a Leica SP6 confocal microscope and images processed in FIJI ImageJ. All settings were kept constant between experimental conditions. For CaLexA experiments, GFP fluorescence was quantified in a region of interest (ROI) based on single optical sections from whole-mount fly VNCs; average GFP signal for each individual sample was calculated by dividing total GFP fluorescence intensity by the number of visible cells (GFP/number of cells).

### RNA sequencing

*Drosophila* VNCs were dissected and prepared for FACS using a protocol adapted from DeSalvo et al., 2014 (*54*). In brief, adult VNCs we dissected in a 30-min window in ice cold PBS (ref: D8537; Sigma-Aldrich), transferred into a dissociation solution of collagenase A (ref: 10103578001; Roche)/PBS with final concentration of collagenase A of 1.5mg/ml, and incubated at 37°C for 30 minutes. Tissues were fragmented by pipetting up/down (10-20 times) and the resulting cell suspension was filtered through a 35µm nylon mesh into a cytometry tube. GFP positive neurons were immediately sorted by flow cytometry (BD FACS Melody) into 5 µl of PBS and flash frozen in liquid Nitrogen. Flash frozen cell samples were sent to the *Cambridge Genomic Services* for library preparation using SMART-Seq® v4 Ultra® Low Input RNA Kit (Takara Bio USA, Inc), followed by RNA sequencing (NextSeq 74 cycle high output 15-20 M reads per sample) and Bioinformatic analysis using Drmelanogaster6 as reference for processing and alignment. Data was extracted as Total read count per gene, read per gene Length (Kp) – RPK, and RPK/million Mapped reads – RPKM. Genes with 0 reads and genes with more than 2-fold difference between biological duplicates were excluded. Lastly, fold-change between samples were calculated as a ratio of experimental samples over wild-type ones (experimental/wild-type).

### Dendrogram generation and homology analysis

We used EMBL-EBI Clustal Omega (http://www.ebi.ac.uk/Tools/msa/clustalw2/) to conduct protein sequence alignment for with the symporter genes. We used the Jalview Version 2-a program (Waterhouse et al., 2009) and the T-Coffee method for multiple sequence alignments and visualisation, and for phylogenetic tree generation. From Jalview we applied JPred 4 programs (*35*) to produce secondary structure prediction. Subsequently, Jmol and Chimera viewers inside the Jalview desktop, and SACS MEMSAT2 (*55*) were employed to generate predicted molecular structures of SLC5 and SLC24 clusters.

### Quantification and statistical analysis

Statistical analyses of data were performed in GraphPad Software Prism using Mann-Whitney U test or one-way ANOVA with the post-hoc Tukey-Kramer test. Error bars in figures represent SEM. Significant values in all figures: ^∗^p < 0.05, ^∗∗^p < 0.01, ^∗∗∗^p < 0.001.

## SUPPLEMENTARY FIGURES

**Fig S1.**
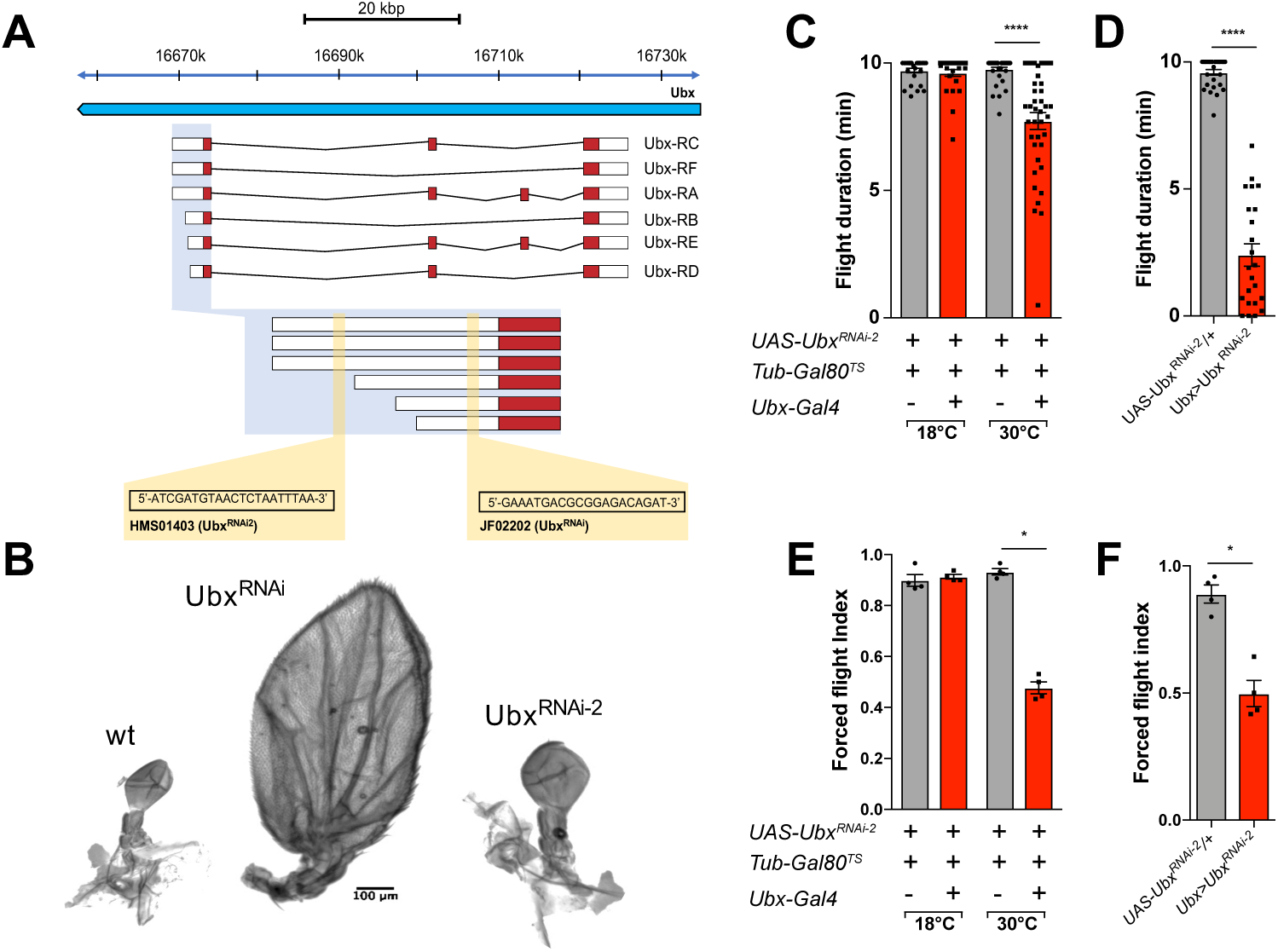
Comparative features of *Ubx^RNAi^* constructs employed. **Related to figure 1.** (**A**) Genomic location of different RNAi targets. (**B**) Effect of different RNAis on haltere morphology. Effect of *Ubx^RNAi-2^* on flight. Average flight duration (**E** and **D**) and forced flight (**E** and **F**) of flies expressing *Ubx^RNAi-2^* in *Ubx* at different conditions. Error bars in figures represent SEM. Significant values in all figures based on Mann-Whitney U test or one-way ANOVA with the post hoc Tukey-Kramer test: ^∗^p < 0.05 and ^∗∗∗^p < 0.001.

**Fig. S2.**
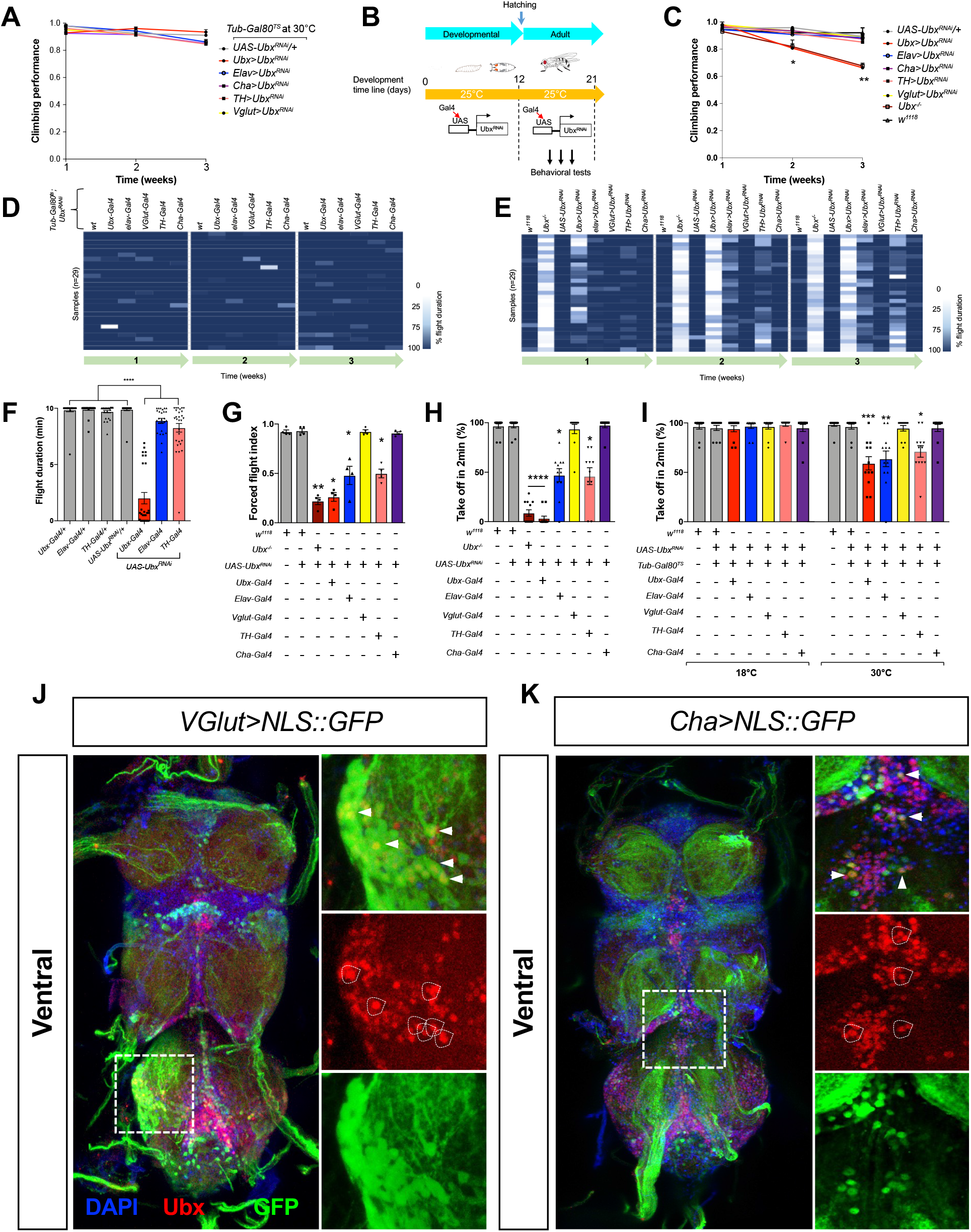
Ubx downregulation affects flight behavior. **Related to figure 1**. (**A**) Climbing performance for conditional neuronal downregulation of *Ubx* in the adult. (**B** and **C**) Strategy and climbing for non-conditional neuronal downregulation of *Ubx* in the adult. (**D**) Flight duration represented as a heat map, for flies expressing *Ubx^RNAi^* in different neuron types under temperature-sensitive *Gal80ts* repressor control, at 18°C. (**E**-**G**) Evaluation of flight performance in non-conditional neuronal downregulation of *Ubx*. Flight duration is represented as a heat map (E) or average (F), and forced flight (G). (**H** and **I**) Take-off ability for non conditional and conditional neuronal downregulation of *Ubx.* See also Movie S1 and S2. Confocal images of the whole ventral nerve cord (VNC) of adult *Drosophila* showing expression of Ubx protein in glutamatergic (**J**) and cholinergic (**K**) neurons. Error bars in figures represent SEM. Significant values in all figures based on Mann-Whitney U test or one-way ANOVA with the post hoc Tukey-Kramer test: ^∗^p < 0.05, ^∗∗^p < 0.01, ^∗∗∗^p < 0.001.

**Fig. S3.**
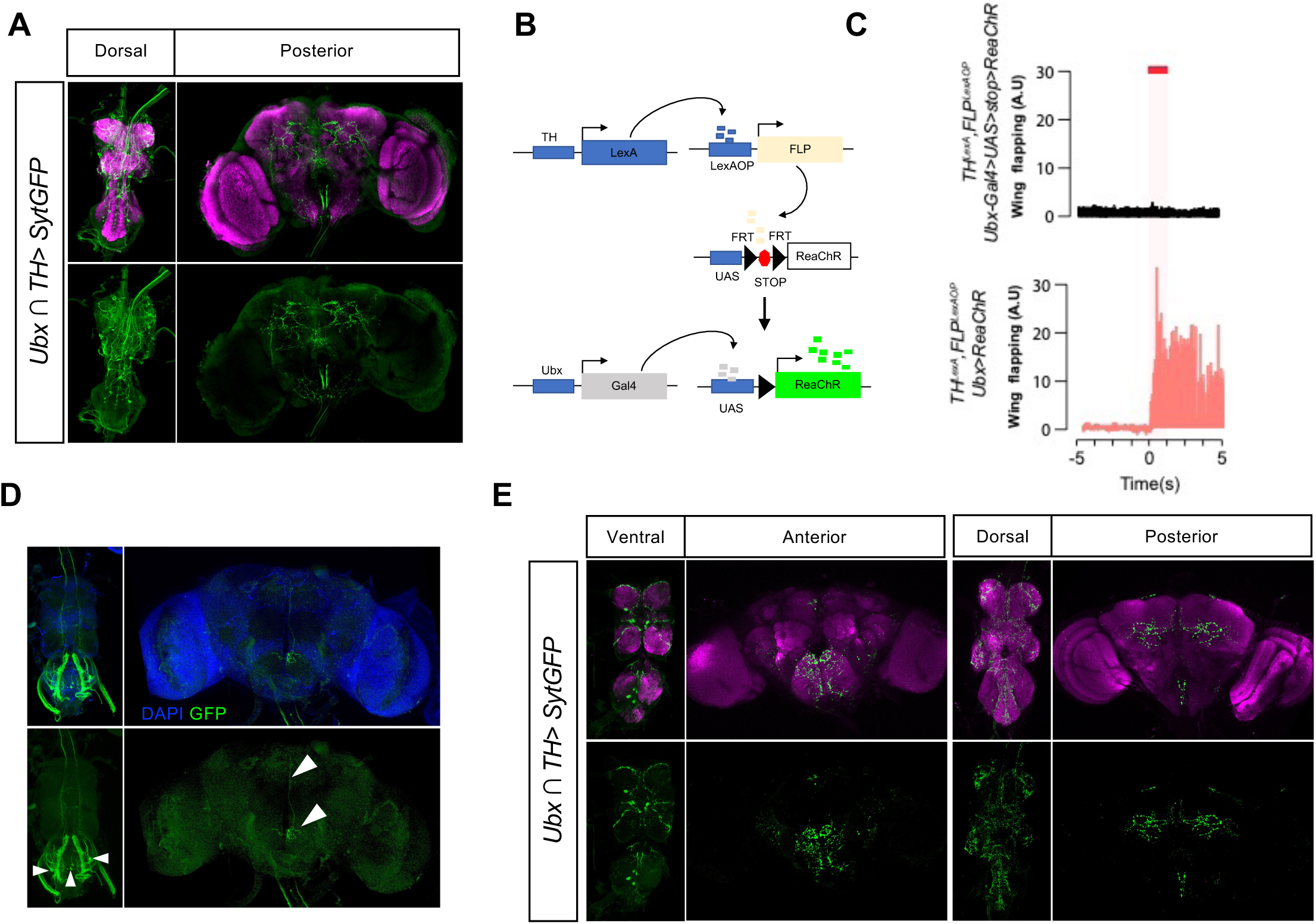
Ubx^+^ TH neurons in the VNC are required for flight. **Related to figure 3**. (**A**) Confocal images of the fly VNC (dorsal view) and brain (posterior view) showing UAS-Myr::GFP expression as driven by Ubx∩TH (UbxGal4.DBD ∩ pleGal4.AD; UAS-Myr:GFP). (**B**-**D**) Activation of Ubx positive TH neurons using FlpOut method. Schematic illustration of the FlpOut method used to activate Ubx^+^ TH neurons (B). Transcription factors TH-LexA and Ubx-Gal4 were combined to drive the operator LexAOP-FLP, and UAS-FRT-stop-FRT-ReaChR. After heat-shock, FRT-mediated removal of a stop codon (red box) leads to expression of channelrhodopsin ReaChR (green boxes) only in Ubx^+^ TH neurons. (**C**) Red light-evoked (red shade) activation of Ubx positive TH neurons expressing channelrhodopsin ReaChR (without stop codon, bottom panel) induces spontaneous flight compared to control (with stop codon, top panel). (**D**) Expression of ReaChR in Ubx positive TH neurons in the VNC (left panel) and brain (right) after light-evoked activation. (**E**) Confocal images of *Drosophila* expressing postsynaptic marker synaptotagmin-GFP (sytGFP) driven by Ubx∩TH (UbxGal4.DBD∩pleGal4.AD; UAS-SytGFP).

**Fig. S4.**
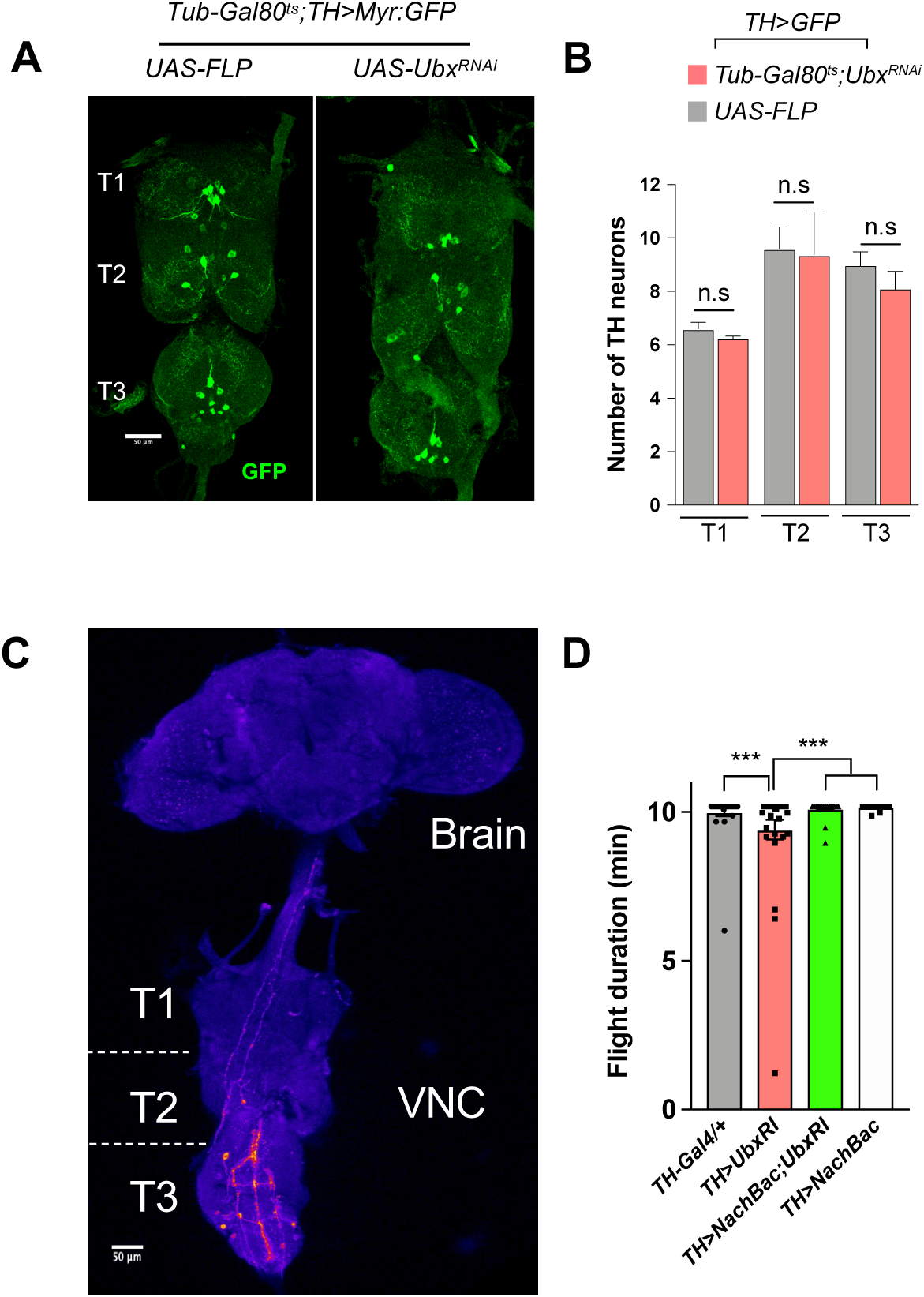
Effects of Ubx downregulation on dopaminergic neuron integrity and activity. **Related to figure 4**. (**A** and **B**) Ubx downregulation in TH neurons does not lead to a reduction in the number of these neurons in VNC thoracic segments (T1, T2, and T3). Genotypes: *TH-Gal4;UAS-Myr:GFP,Tub-Gal80ts;UAS-FLP* and *TH-Gal4;UAS-Myr:GFP,Tub-Gal80ts;UAS-Ubx^RNAi^*. (**C**) Flight induces an increase in the CaLexA signal in the TH neurons of VNC and not in the brain. (**D**) non conditional expressing of NaChBac in TH neurons rescues the flight deficit induced by Ubx downregulation. Error bars in figures represent SEM. Significant values in all figures based on Mann-Whitney U test or one-way ANOVA with the post hoc Tukey-Kramer test: ^∗∗∗^p < 0.001.

**Fig. S5.**
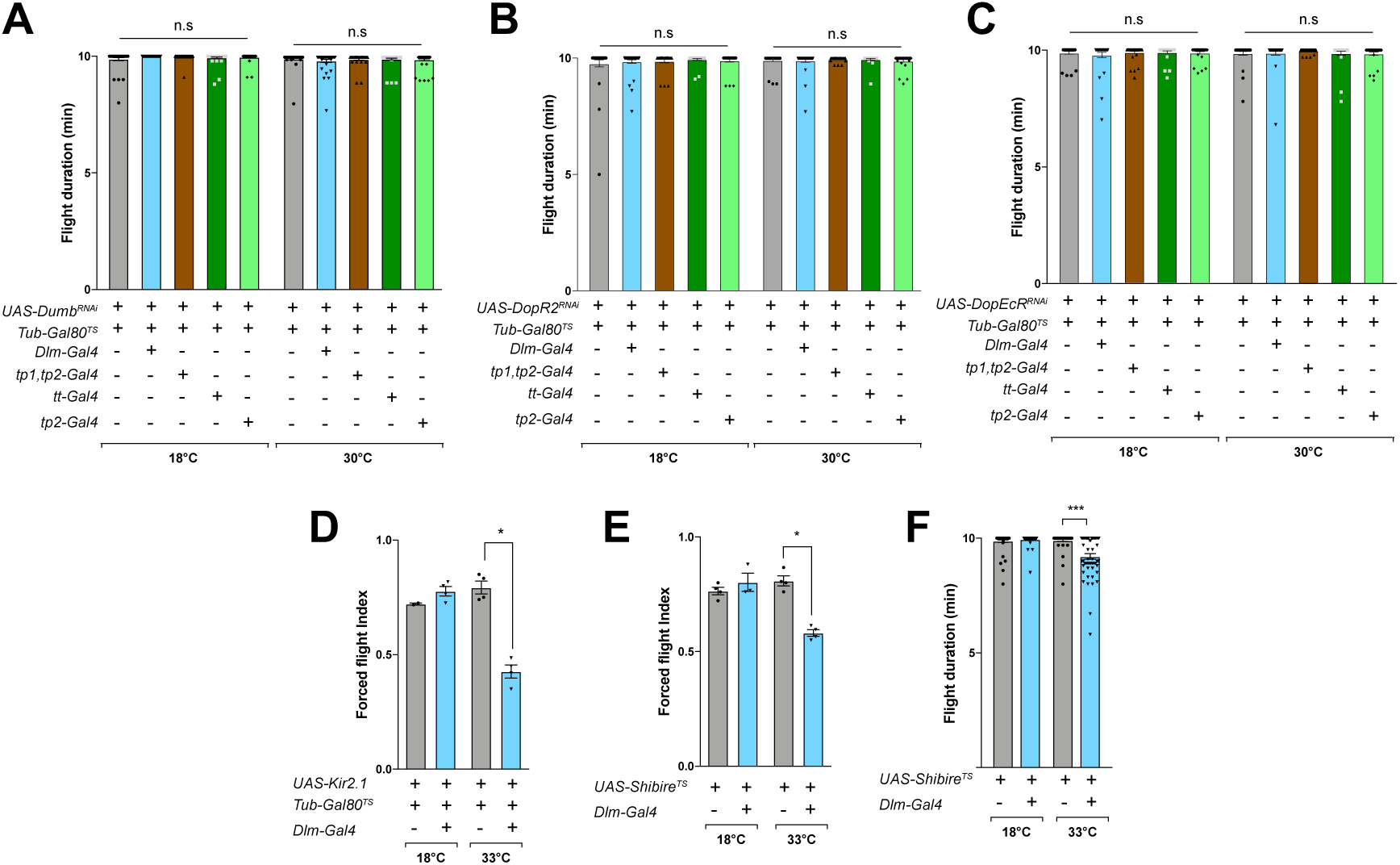
A subset of flight motor neurons conveys TH neuron signals. **Related to figure 5.** (**A** and **B**) Evaluation of flight performance under conditional neuronal downregulation of dopaminergic receptors: Dop1R1/Dumb (A), DopR2 (B), or DopEcR (C). (**D**) Reduction in the activity of Dlm neurons using Kir2.1 leads to a flight deficient phenotype (**E** and **F**) Expression of *Shibire^ts^* in Dlm neurons reduces average duration of flight and forced flight. Error bars in figures represent SEM. Significant values in all figures based on Mann-Whitney U test or one-way ANOVA with the post hoc Tukey-Kramer test: ^∗^p < 0.05, ^∗∗∗^p < 0.001.

**Fig. S6.**
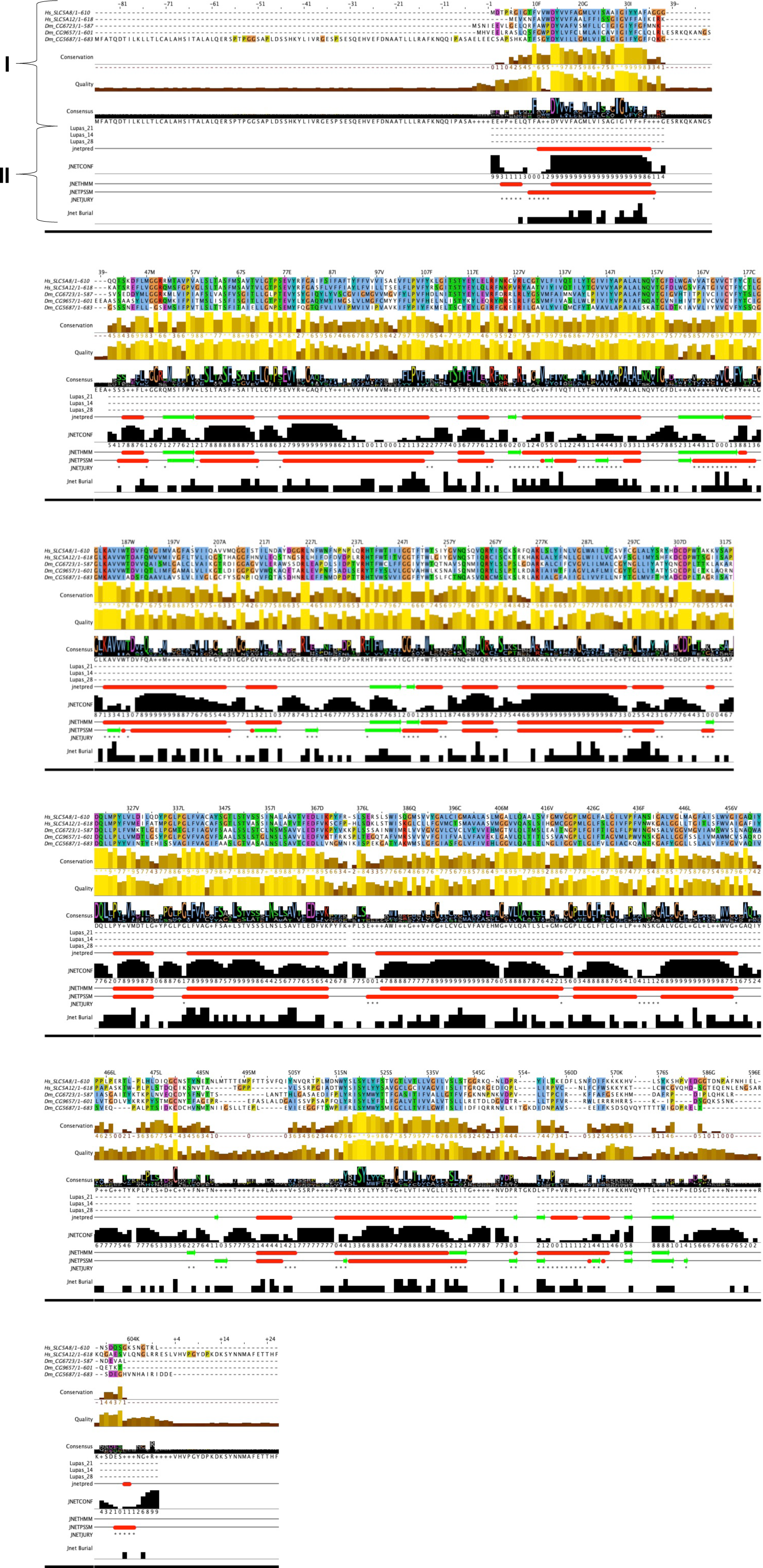
Structural alignment of *Drosophila* (Dm) SLC genes to human (hs) SLC5A12 and SLC5A8 genes. **Related to figure 6.** Part I represents a multiple sequence alignment made with the Jalview program. A colour code represents conservation of distinct aminoacid residues in the alignment. Blue:hydrophobic, red:positive charge, magenta:negative charge, green:polar, pink:cysteines, orange:glycines, yellow:prolines, cyan:aromatic. Part II, JPred secondary structure prediction. Helices are marked as red tubes, and sheets as dark green arrows JNetPRED: consensus prediction. JNetCONF: confidence estimates for the prediction. High values mean high confidence. Lupas_21, Lupas_14, Lupas_28: Coiled-coil predictions for the sequence. Jnet Burial: Prediction of Solvent Accessibility. JNetALIGN: Alignment based prediction. JNetHMM: HMM profile-based prediction. JNETPSSM: PSSM based prediction. A ’*’ in this annotation indicates that JNETJURY was invoked to rationalise discrepant primary predictions.

**Fig. S7.**
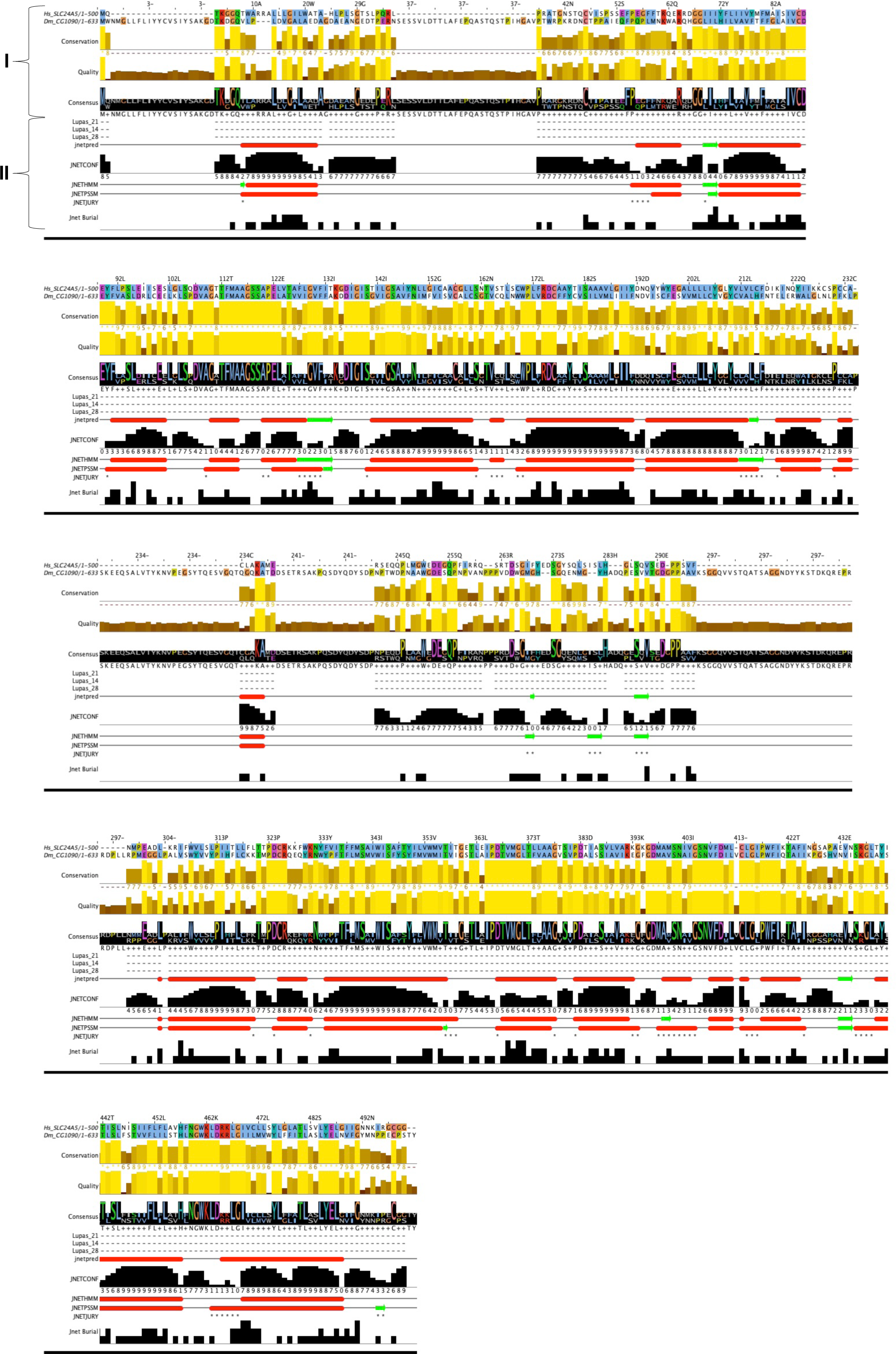
Structural alignment and prediction of Drosophila SLC genes orthologous to human SLC24A5. **Related to figure 6.** Part I represents a multiple sequence alignment made with the Jalview program. A colour code represents conservation of distinct aminoacid residues in the alignment. Blue:hydrophobic, red:positive charge, magenta:negative charge, green:polar, pink:cysteines, orange:glycines, yellow:prolines, cyan:aromatic. Part II, JPred secondary structure prediction. Helices are marked as red tubes, and sheets as dark green arrows JNetPRED: consensus prediction. JNetCONF: confidence estimates for the prediction. High values mean high confidence. Lupas_21, Lupas_14, Lupas_28: Coiled-coil predictions for the sequence. Jnet Burial: Prediction of Solvent Accessibility. JNetALIGN: Alignment based prediction. JNetHMM: HMM profile-based prediction. JNETPSSM: PSSM based prediction. A ’*’ in this annotation indicates that the JNETJURY was invoked to rationalise significantly different primary predictions.

**Fig. S8.**
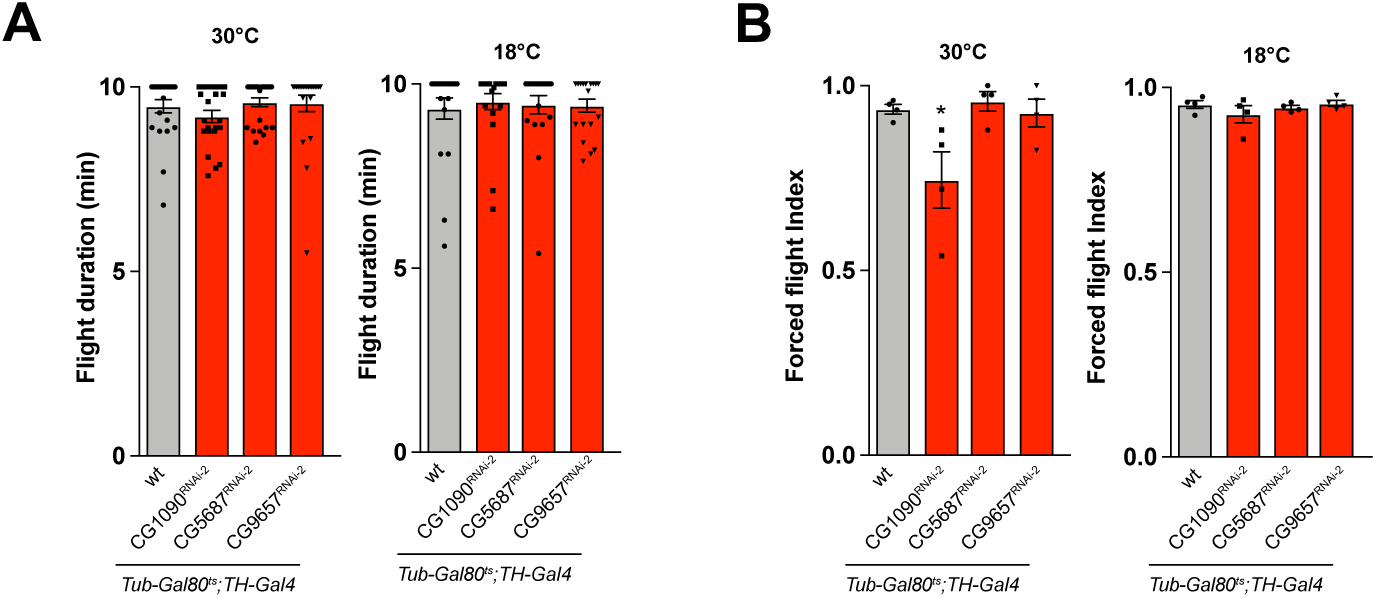
Conditional downregulation of symporter genes in TH neurons affect flight behaviour. **Related to figure 6.** Behavioural effects of conditional downregulation of symporter genes by means of alternative RNA interference constructs. (**A**) Flight maintenance. (**B**) Forced flight.

